# Unraveling dynamically-encoded latent transcriptomic patterns in pancreatic cancer cells by topic modelling

**DOI:** 10.1101/2023.03.11.532182

**Authors:** Yichen Zhang, Mohammadali (Sam) Khalilitousi, Yongjin P Park

## Abstract

Building a comprehensive topic model has become an important research tool in single-cell genomics. With a topic model, we can decompose and ascertain distinctive cell topics shared across multiple cells, and the gene programs implicated by each topic can later serve as a predictive model in translational studies. Here, we present a Bayesian topic model that can uncover short-term RNA velocity patterns from a plethora of spliced and unspliced single-cell RNA-seq counts. We showed that modelling both types of RNA counts can improve robustness in statistical estimation and reveal new aspects of dynamic changes that can be missed in static analysis. We showcase that our modelling framework can be used to identify statistically-significant dynamic gene programs in pancreatic cancer data. Our results discovered that seven dynamic gene programs (topics) are highly correlated with cancer prognosis and generally enrich immune cell types and pathways.

## Introduction

Single-cell RNA-seq technology has been successfully applied to profile regulatory-genomic changes in studying many human disease mechanisms. Our capability to measure single-cell-level mRNA molecules has dramatically changed our research paradigm in genomics and translational medicine. A typical single-cell study implicitly assumes observed transcript levels as a static value, considering that every cell is fixed at a particular state. Recently, researchers have developed a complementary method to measure gene expression dynamics (the speed of splicing) by taking the difference between the spliced and unspliced counts in scRNA-seq profiles.^1^ Several methods have extended the original method pioneered by La Manno and coworkers. Notably, scVelo method generalizes to recover gene-level ordinary differential equations (ODEs), allowing each gene to take independent time scales.^2^

### Why is it difficult to estimate full-scale dynamics in data sets with limited snapshots?

However, probabilistic inference of full-scale dynamics often poses a substantial challenge, and the inferred rate parameters may greatly vary depending on the normalization and embedding methods.^3^ Although a newly-developed machine learning based on a mixture of ODE models improved the robustness and accuracy in single-cell data profiled in developmental processes,^4^ existing velocity analysis methods rely on a critical assumption unmet by most single-cell data sets at a study design level. Most single-cell datasets, especially those collected from patient-derived cancer samples, only span over several snapshots of full developmental, evolutionary or disease progression processes. In human case-control studies, cells may not have reached steady states in the disease progression process and are likely to fail to provide enough information for most genes and pathways. Such discontinuity and sparsity in data collection somewhat force statistical inference algorithms to rely on an unrealistic steady-state assumption and interpolated data points with high uncertainty.^3,5^

### Why do we need a topic model for transcription dynamics?

Nevertheless, gene expression dynamics implicated by the transcript-level difference between the spliced and unspliced counts provide a valuable perspective in single-cell data analysis, making single-cell analysis more valuable beyond conventional static analysis. To overcome the limitations of poor and incompleteness in single-cell RNA velocity analysis, we propose a new modelling framework, DeltaTopic, short for Dynamically-Encoded Latent Transcriptomic pattern Analysis by Topic Modelling. DeltaTopic combines two ideas: (1) latent topic analysis that will guide unsupervised machine learning for discovering new dynamic cell states, (2) application of first-order approximation to learn robust relationships between the spliced and unspliced counts instead of estimating a full trajectory of ODE models. For a latent topic model, we view each cell as a document and each gene as a word to make model parameters directly interpretable while keeping the Bayesian model’s capability to impute missing information. The simplified dynamic model also permits an intuitive interpretation of spliced-unspliced differences as multiplicative “delta” parameters in the model.

We developed and applied our DeltaTopic approach to single-cell datasets on pancreatic ductal adenocarcinoma (PDAC), one of the most challenging cancer types with a poor prognosis. In the latent space, our model identified cancer survival-specific topics marked by a unique set of gene expression dynamics. We also find DeltaTopic further dissected sub-topics clumped together in traditional clustering methods implicating novel gene modules and cell states that are dynamically controlled along with the cancer progressions.

## Results

### Single-cell transcriptomic dynamics in Pancreatic Ductal Adenocarinoma (PDAC) studies

We preprocessed single-cell expression data sets available in two large-scale, multi-individual studies.^6,7^ We extracted both spliced and unspliced count vectors from the original short-read sequencing files for each cell, using Kallisto^8^ and UMI-BUS^9^ tools. We consolidated all the samples/batches into one file set using our customized utility functions `rcpp_mmutil_merge_file_sets` available in `mmutilR` library (https://causalpathlab.github.io/mmutilR). Applying cell-level quality control steps, which filtered out cells with too few counts and high mitochondrial gene expression activities, we retained 227,331 (91.13%), discarding 22,126 cells (8.87%) by applying these Q/C steps out of 249,457 cells. The quantification algorithm results in two types of gene expression vectors for each cell–one for the spliced and the other for the unspliced counts. We measured 22,836 features (the spliced and unspliced genes) on a total of 227,331 cells, and only 329,824,833 elements were non-zero (6.35%).

Overall, we have two types of high-dimensional sparse matrices as input data on total *G*=11,418 genes across *N*=227,311 cells: (1) 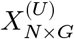 for the unspliced counts and (2) 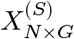 for the spliced transcript counts. Our goal is to identify latent factors/topics and the corresponding topic-specific gene expression frequency parameters–one for the unspliced and the other for the spliced components. No additional preprocessing steps, such as gene selection, batch adjustment, selection of principal components, and data transformation, were necessary for topic modelling. Since the underlying multinomial likelihood is less affected by potential effects of unwanted stochasticity than other probability models.^10,11^

### Overview of our approach

#### A Bayesian approach to identify sparse cell topics in scRNA-seq data (BALSAM)

We developed a Bayesian topic modelling approach, extending the Embedded Topic Model framework^12^ with elementwise spike-and-slab prior probability.^13^ Our BALSAM approach, short for Bayesian Latent topic analysis with Sparse Association Matrix, views cells as an admixture of gene topics to summarize static transcriptome patterns from raw gene expression count data (Fig 1A). BALSAM relies on Variational AutoEncoder (VAE) to learn cell topics and infer the gell-topic relationship. The encoder transforms the expression space to a latent topic space through a stack of non-linear layers (NN1), outputting a vector of relative topic proportions for each cell. The decoder generates a dictionary matrix *β* from a sparse-inducing prior called spike-slab to ensure only a small number of genes are selected for each topic. The resulting dictionary matrix *β* is passed to a generalized linear model (GLM) along with topic proportion *θ* to estimate normalized gene frequency *λ*. Using *λ* as the parameter, we compute the likelihood of the expected gene count from a multinomial distribution. We provide detailed descriptions of the BALSAM, sparse priors, and Variational inference algorithms in Methods.

**Figure 1.**
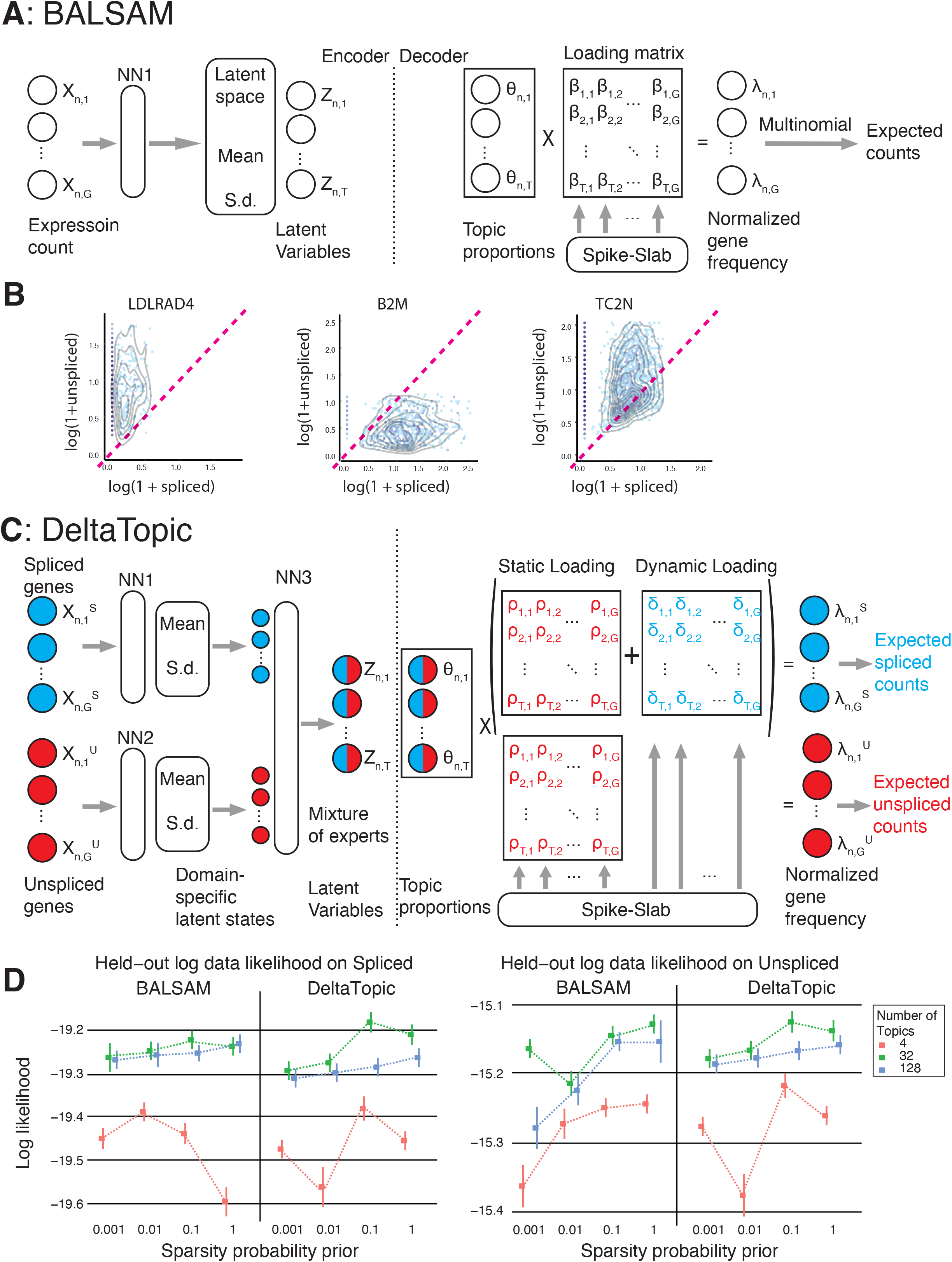
Modelling single-cell transcription dynamics with deep probabilistic topic models. **(A)** BALSAM: Given as input a raw gene expression count matrix, Balsam learns cell topics to represent cell types or cell states using neural networks. The encoder transforms the expression space to a latent topic space through a stack of non-linear layers (NN1). The decoder generates biologically interpretable topic embeddings for each topic from a sparse-inducing prior (spike-slab), and estimates the normalized gene frequency *λ* from topic proportion *θ* and loading matrix *ρ* by a generalized linear model (GLM). Using *λ* as the parameter, we compute the likelihood of the expected gene count from a multinomial distribution. **(B)** Scatterplots of top genes in 3 disease-relevant delta topics, with 2-D density curves imposed. Y-axis: unspliced gene counts (log-transformed), X-axis: spliced gene count ((log-transformed)). The red dashed line indicates steady state, where the spliced and unspliced gene is in the same amount. **(C)** DeltaTopic: Given as input a pair of spliced and unspliced gene expression count matrices, DeltaTopic transforms each to a latent space through two independent Balsam encoders (NN1, NN2) and combines the information to form a shared latent space through a fusion layer (NN3). The decoder of DeltaTopic generates two loading matrices from sparse-inducing spike-slab priors to decompose spliced and unspliced transcription patterns into static and dynamic topics. The static topic loading matrix *ρ* serves as a background loading for both spliced and unspliced genes. For spliced genes, the dynamic topic loading matrix is added to the static loading matrix to learn transcriptome dynamics. **(D)** Model evaluation on held-out data likelihood (spliced and unspliced). Y-axis: the average held-out data likelihood and 95% confidence interval. X-axis: sparsity probability prior.

#### Dynamically-Encoded Latent Transcriptomic pattern Analysis by Topic modelling (DeltaTopic)

In designing our DeltaTopic approach, we were inspired by common patterns that we repeatedly observed in gene-level data (Fig. 1B):

- **Sparsity due to technologies**: Uncontrollable drop-out events may assign a substantial fraction of gene expression counts to a zero value, obfuscating our delineation between the undetectable and unexpressed genes. For instance, the spliced counts of the *LDLRAD4* gene are essentially zero in many cells, even whilst the unspliced counts are positive (Fig. 1B on the left). Similarly, an excess of zero values in the unspliced was observed in *B2M* (Fig. 1B in the middle); for the *TC2N* case, both sides contain many zero values (Fig. 1B on the right).
- **Sampling bias in a temporal axis**: In contrast to what an analytical ODE solution predicts, we tend to have only a limited portion of the whole phase diagram of RNA velocity.^1^ Parametric inference for an ODE model would remain unidentifiable, suggesting multiple similarly likely solutions without a strong steady-state assumption. Again, as can be seen in the exemplary genes (Fig. 1B), genes were differentially regulated in the same data set, potentially suggesting the existence of multiple modes in transcription dynamics.

At least in our cancer study, we found almost all genes fail to reach steady states showing a characteristic phase diagram.^1^ Therefore, we focused on modelling short-term directional information implicated by RNA velocity in an intuitive model regressing the spliced on the unspliced count data, gene by gene and topic by topic. DeltaTopic model extends the BALSAM model to ascertain common cellular topic space and topic-specific relationships between the unspliced and spliced data (Fig. 1C). Since a generalized linear model framework provides a flexible way to capture relationships across many data modalities, our approach can be easily extended to other similar tasks.

Given a pair of spliced and unspliced gene expression count matrices as input, DeltaTopic transforms each to a latent space through two independent BALSAM encoders (NN1, NN2). The latent variables with encoded information from two encoders were combined by taking their average^14,15^ to obtain a shared topic vector as a mixture of experts from the spliced and unspliced latent space (Fig. 1C, fusion layer). The decoder then generates two dictionary matrices from sparsity-inducing spike-slab priors to decompose spliced and unspliced transcription patterns into static and dynamic topics. The static topic dictionary matrix *ρ* sets a background level for both spliced and unspliced genes. The dynamic topic dictionary matrix *δ* for spliced genes determines the directionality–activated vs. inhibited–compared with the background dictionary matrix *ρ* at a topic level. The decoder model generates the normalized frequency for the spliced and unspliced counts as a linear combination of multiple dynamic/static topics weighted by cell-level topic proportions. We implemented the model in `PyTorch` and performed posterior inference using NVIDIA RTX 3080 GPU.

#### Training deep learning models and generalization performance

To evaluate each model’s generalization performance, we tested the likelihood of data that were held out during training for BALSAM and DeltaTopic. We randomly split the cells by a 9-to-1 ratio to form training and test sets. For BALSAM, we trained two independent models, one with the spliced gene count as input and the other with the unspliced gene count. We validated the held-out log data likelihood in their corresponding domain (e.g., spliced or unspliced). For DeltaTopic, we trained a unified model with the pairs of spliced and unspliced counts as input and validated the held-out log-likelihood in both domains. To study the effects of the latent dimension and the sparsity probability prior on the model generalizability, we trained BALSAM and DeltaTopic with different numbers of latent topics (4, 32, 128) and set sparsity probability prior to *π* = 0.001,0.01,0.1,1. Each training was repeated five times with different random seeds.

We found the DeltaTopic with 32 topics and sparsity probability prior *π* = 0.1 is overall the best-performing model, compared to other choices of latent dimension and sparsity level (Fig. 1D). While for BALSAM; the best-performing model is the one with 32 topics and sparsity probability prior *π* = 0.01 in the spliced domain; and *π* = 1 in the unspliced domain. With this choice of hyperparameters, DeltaTopic yields better hold-out log-likelihood scores than BALSAM in both types-the spliced and unspliced data, suggesting that modelling the relationships between two related data types can improve robustness. For both models, finding the right model complexity (the number of topics) was necessary. Although a more fine-grained hyperparameter search, empowered by a Bayesian optimization method, is desired, our results suggest that underfitting with four topics and overfitting with 128 topics tend to yield inferior generalization performance. A wider error bar is observed in the underfitting model (with four topics), suggesting the model is more sensitive to the choice of initialization. In general, Bayesian sparsity priors avoid overfitting issues since we observed that a model with either 32 or 128 topics performs similarly regarding the hold-out log-likelihood scores in both data types. Indeed, using a high number of topics and stringent sparsity hyperparameters enforced, our Bayesian framework can automatically determine the right balance between bias and variance, shutting off unnecessary topics and leading to slightly improved generalization performance.

#### DeltaTopic provides novel insights into pancreatic cancer etiology

Having an optimal configuration of hyperparameters established, we trained our topic models on the full data set. We then asked whether the genes/features discovered by DeltaTopic are novel and different from the genes found by the topic models that were trained on either the spliced or unspliced counts alone. In order to answer the question, we linked the delta topics to the counterparts found by BALSAM. Using the cell-level topic proportion estimates, namely *θ_i_*, per each model, almost all the topics found in the Delta Topic were connected to the ones found in BALSAM (Supplementrary Fig. 1). For instance, Delta topic #4 and BALSAM topic #4, Delta topic #10 and BALSAM topic #8, Delta topic #11 and BALSAM topic #21 are highly overlapping in their membership. However, topics with less than 2% of the total population cells were not included for further investigation for brevity. DeltaTopic marginally improves the resolution in cell clustering, providing a finer view of transcriptome dynamics. Some topics found by BALSAM can be portioned into two or more Delta topics. For instance, BALSAM #32 can be dissected into two Delta topics, #3 and #7; likewise, BALSAM # 27 can be divided into three Delta topics, #6, #14, and #30 (Supplementrary Fig. 1).

To better understand the activities of the top genes prioritized by the directional/velocity information (*δ_t_*) in each topic significantly differ from *ρ_t_* and *β_t_*, we investigate therapeutic impacts of the three different gene sets. We estimated the topic-specific risk scores for pancreatic cancer samples available in ICGC Data Portal and correlated the estimated risk scores with the survival outcomes of each individual. Bulk gene expression data sets for three independent pancreatic cancer studies are publicly available with donor-level information in the ICGC Data Portal (https://dcc.icgc.org/releases/current/Projects), including pancreatic cancer samples in the Canadian cohort (N=234 donors on unique 53,800 transcripts), Australian cohort (N=91 donors on unique 42,346 transcripts) and the US cohort (N=142 donors on unique 20,009 genes). We computed donor-level scores by taking average gene expression values weighted by topic-specific gene frequency vectors (vectors in the dictionary matrices): 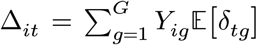. Similarly, we can estimate the other two types of topic-specific scores by 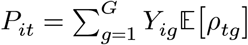, and 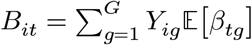.

We correlated these topic and model-specific estimated risk scores with survival outcomes (time-to-event) using a regularized Cox proportional hazard regression model. To account for cohort-specific bias in the survival data, we conducted meta-analysis, aggregating the hazard ratio statistics independently estimated in each cohort by an inverse variance-weighted (IVW) average approach. The IVW approach prioritizes topics found significant consistently across three cohorts while penalizing statistically significant topics only in one or two cohorts. Denoting 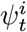 be hazard ratio estimate for topic *t* and cohort *i*, and 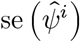 be the standard error. For each topic *t*, we can obtain a summary hazard ratio estimate 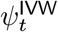 by aggregating cohort-specific hazard ratio 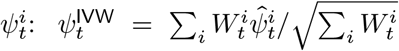, where 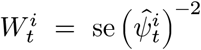, and computed the p-value by 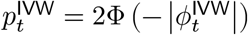.

In Fig. 2B, we plotted each topic’s hazard ratio estimates and p-values. Topics with p-value smaller than 0.05 and absolute hazard ratio estimate greater than 1.5 × *e*^-3^ are interpreted as “survival-relevant,” coloured in red and blue for up-regulating and down-regulating topics, respectively. 7 survival-relevant topics were identified using *δ_t_*, whereas none and only one topic was identified using *ρ_t_* and *β_t_*. Among seven survival-relevant delta topics, topics 11, 4, 29, and 7 were up-regulating survival topics correlated inversely with hazard ratio estimates. Topic 26, 6, and 19 were down-regulating survival topics.

**Figure 2.**
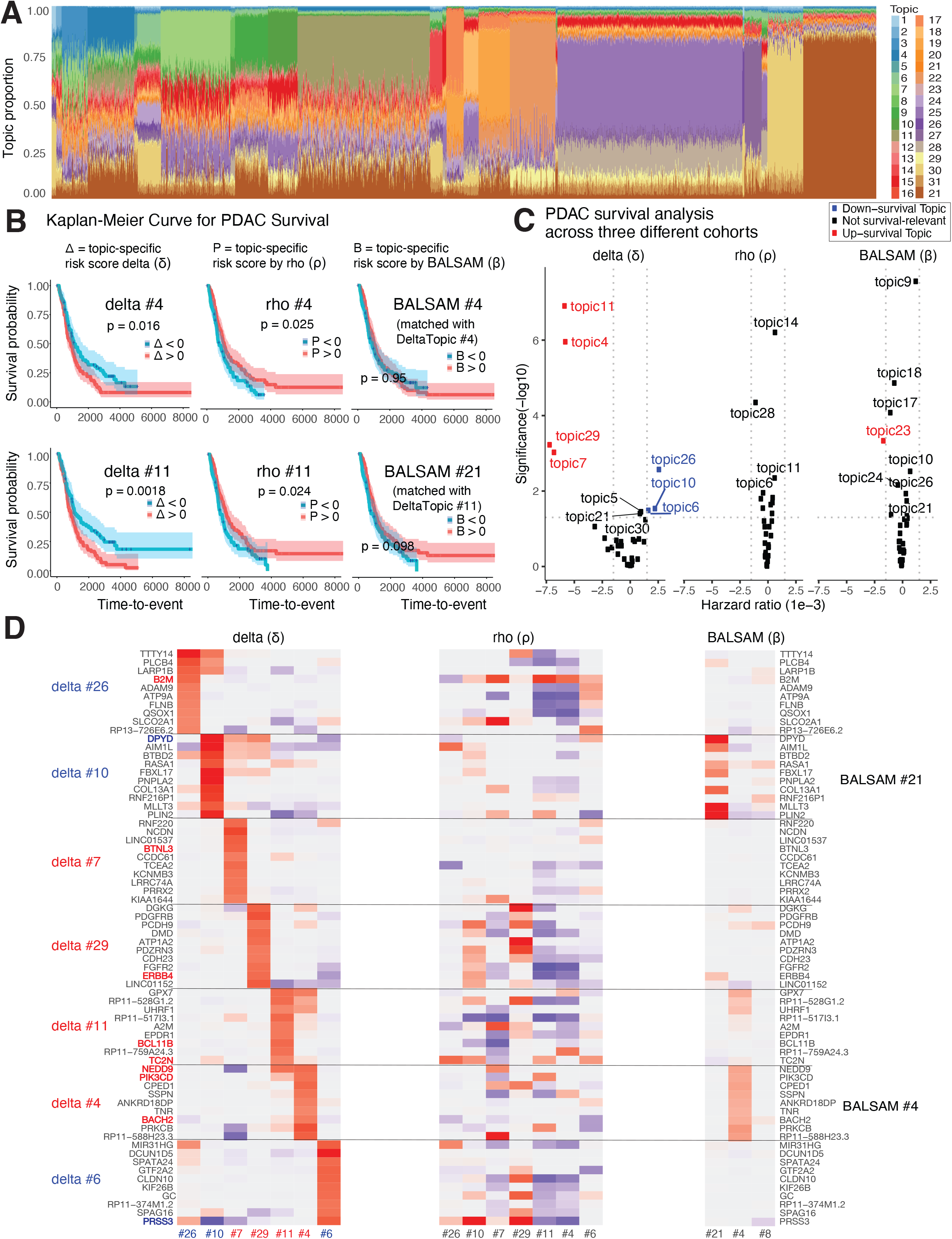
DeltaTopic approach identifies disease-relevant cell topics, implicating putative causative regulatory programs. **(A)** Relative proportion of 32 cell topics on 227,331 cell from PDAC study estimated by the DeltaTopic model. **(B)** Kaplan-Meier survival curves for individuals from the TCGA-PDAC-CA cohort with postive and negative topic-specific risk scores. P-values are computed by log-rank test comparing positive and negative risk groups in survival probability. **(C)** Volcano plot showing hazard ratio and p-value for the association between pancreatic cancer survivals and cell topic proportions from three cancer studies, i.e., TCGA-PDAC-US, TCGA-PDAC-CA, and TCGA-PDAC-AU. Each point represents an aggregated hazard ratio measure and p-value from the three cohorts by meta-analysis. Y-axis: p-value after −log10 transformation, the more significant point appears on the top. X-axis: the hazard-ratio estimate from the Cox proportional hazard model. Survival-relevant cell topics are coloured red and blue for up- and down-regulation in PDAC survival. The two vertical dashed lines correspond hazard ratio cutoff at ±1.5 × *e*^-3^. The horizontal line is the p-value cutoff at 0.05.

Not all topics are directly comparable across different models. For some of them, we were able to compare those topics across different models if they are paired (Supplementary Fig. 1A). Of those matched, we found many cases where the scores derived from Delta Topic are significantly associated with the survival outcome whereas the scores derived from the unimodal topic model are not strongly associated. For instance, topic 4 in DeltaTopic and topic 4 in BALSAM predict the disease prognosis quite differently. Two patient groups stratified by the DeltaTopic (Δ) 4 follow significantly different disease prognoses (p=0.016). In the same way, the delta topic score 11 can partition the cohort into two distinctive subgroups in their prognosis (p=0.0018).

For the seven delta topics significantly associated with the survival outcome, topics 11, 4, 29, 26, 10, 7, and 6 (Fig 1C), we further investigated their top genes/features that define the characteristics of their topics, meaning strong |*δ_tg_*| values. Interestingly, these anchor genes show highly topic-specific activities (Fig. 2D, left). However, the same set of genes showed weaker topic-specific patterns (Fig. 2D, middle) and in the BALSAM model (Fig. 2D, right). Our results suggest that our DeltaTopic approach can distinguish genes that are differentially regulated in transcriptomic dynamics from differentially expressed at the static level, and they can play a pivotal role in cancer progression and metastasis.

Interestingly, the top genes in survival-relevant delta topics were related to PDAC and other cancer-related processes. *B2M*, among the top genes in the down-survival topic (delta #26) were associated with poor prognosis in PDAC.^16^ *DPYD*, among the top genes in another down-survival topic (delta #10), was found to be overexpressed in the PDAC sample with a poor prognosis in the immunohistochemical analysis.^18^ Furthermore, *NEDD9*, selected for delta #4 and #11, is a prognostic maker in pancreatic cancer progression.^19^ *ERBB4* in topic 29 is known to accelerate PDAC development and progression.^20^ These top genes were also related to other cancer cell processes. *BCL11B*, in up-survival topic delta #11, is a well-established tumour suppressor gene in lymphoma and leukemia.^21^ Another oncogene *TC2N* was also found in the same delta #11, suggesting its role in helping tumour cells survive by suppressing the p53 signalling pathway.^22^ *BTN3A* in topic #7 can be regulated by several signals induced by cancer cell or their microenvironment^23^.

#### DeltaTopic model also identifies disease-relevant cell types and pathways

We next evaluated the extent to which the cell topics inferred by DeltaTopic reflect static and dynamic transcriptome patterns. In general, we found rho topic has high correspondence to cell type differentiation (Fig. 3A), while the delta topic loadings are more relevant to gene activities (Fig. 3 B1, B2, B3). Our results suggest that the *ρ* topics are suitable to capture cell-type-specific marker genes and recapitulated known immune cells, such as B-cell, T-cell and macrophages, endocrine and endothelial cells, and multiple subtypes of ductal cells. Topics 15, 18, 25, and 28 correspond to one subtype of the ductal cells, and the other topics 14, 9, 27, 30, 26, and 6 are specifically matched with the second ductal cells found in the original study.^6^ Topic 4 and 11 are likely associated with immune activities, as they mostly constitute immune cells’ activities.

**Figure 3.**
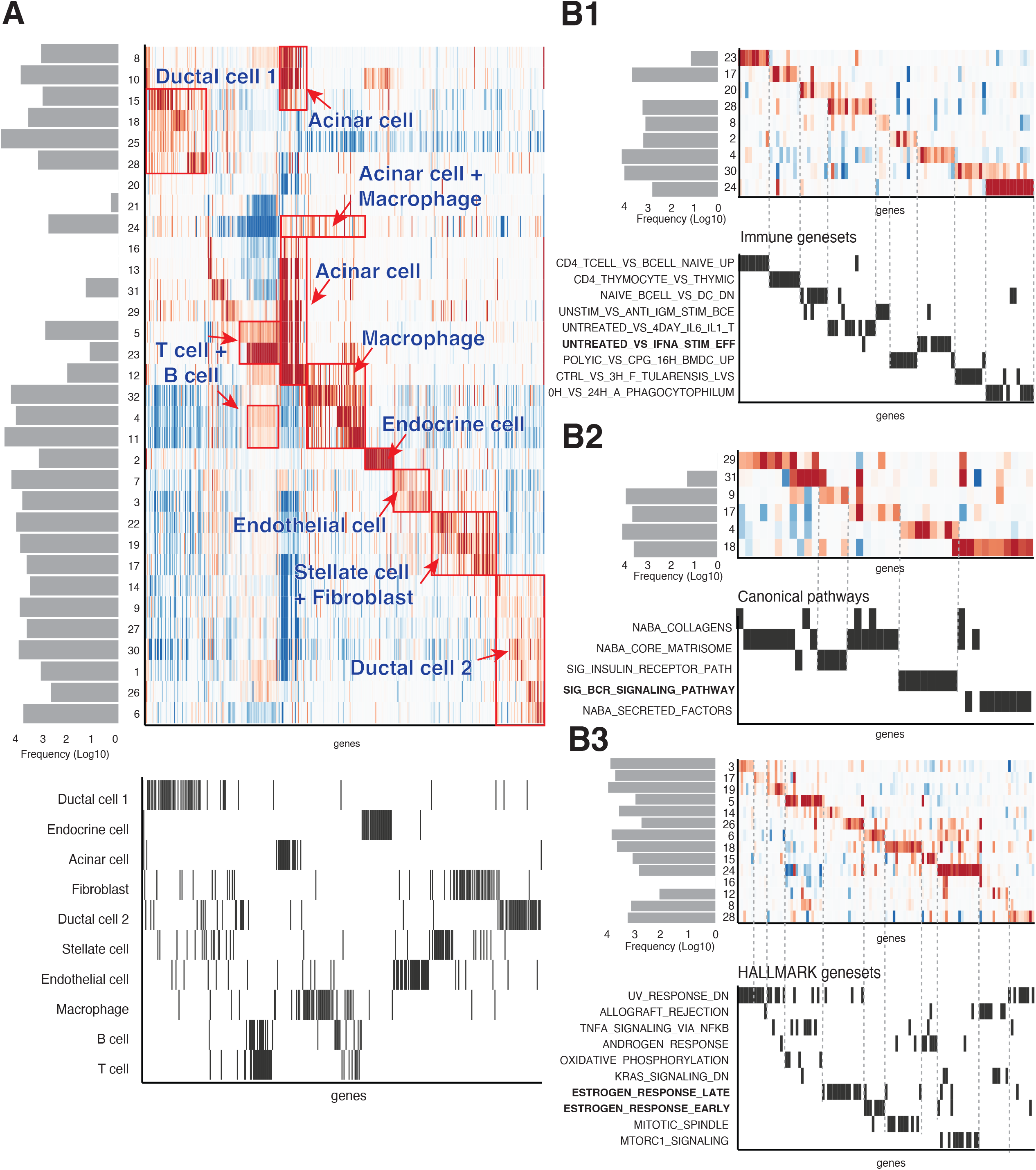
DeltaTopic approach uncover both static and dymanic transcriptome patterns. **(A)** Heatmap of static loading matrix with topic sizes visualized as a barplot on the left. On the bottom, the markers gene of 10 cell types from [Peng et al.]. **(B)** Gene set enrichment analysis of dynamic loading matrix. **B1**: ImmuneSig gene sets; **B2**: KEGG gene sets; **B3**: Hallmark gene sets. All three gene sets are from MsigDB database (https://www.gsea-msigdb.org/gsea/msigdb/). Only significant genesets and their corresponding cell topics are shown.

Gene set enrichment analysis of the Molecular Signatures Database^24–26^ using `fgsea` package^27^ provided another line of evidence. We observed their enrichment with the gene set “CD4 TCELL VS BCELL DN”, which comprises genes differentially regulated between T and B cells. Unlike the static dictionary matrix *ρ*, the dynamic one *δ* is more likely to capture the gene expression changes more pertinent to immune mechanisms. For instance, Topic 4 enriches two immune gene sets–“UNTREATED VS IFNA STIM CD8 TCELL 90MIN UP” and “SIG BCR SIGNALING PATHWAY.” Both of them were characterized to control tumour growth mechanisms.^28^ Other topics 6 and 26 also bear many cancer-regulatory genes, constituting “ESTROGEN RESPONSE EARLY” and “ESTROGEN RESPONSE LATE” pathways, also well-aligned with the previous study,^29^ reporting high expression of estrogen receptor beta genes in pancreatic adenocarcinoma samples resected from a poor prognosis group.

#### Vector fields reconstructed from DeltaTopic dictionary visualize distinctive disease trajectories

We projected the estimated transcriptome dynamics (velocities) onto a 2D space constructed by its eigenvectors (Fig. 4A). Each headed arrow represents the estimated velocities for a cell, with the starting point for unspliced genes and the ending point for its spliced counterpart. We simply carried out the projection as follows: (1) Perform singular value decomposition (SVD) on *ρ* dictionary matrix (with rank 2) *ρ* = *UDV^T^* to get its 2D representation *V*, (2) project *ρ* + *δ* onto the same eigenspace for spliced genes by *V*^spliced^ = (*ρ* + *δ*)*UD*^-1^, (3) and connect *V* to *V*^spliced^ for each cell to visualize velocities.

**Figure 4.**
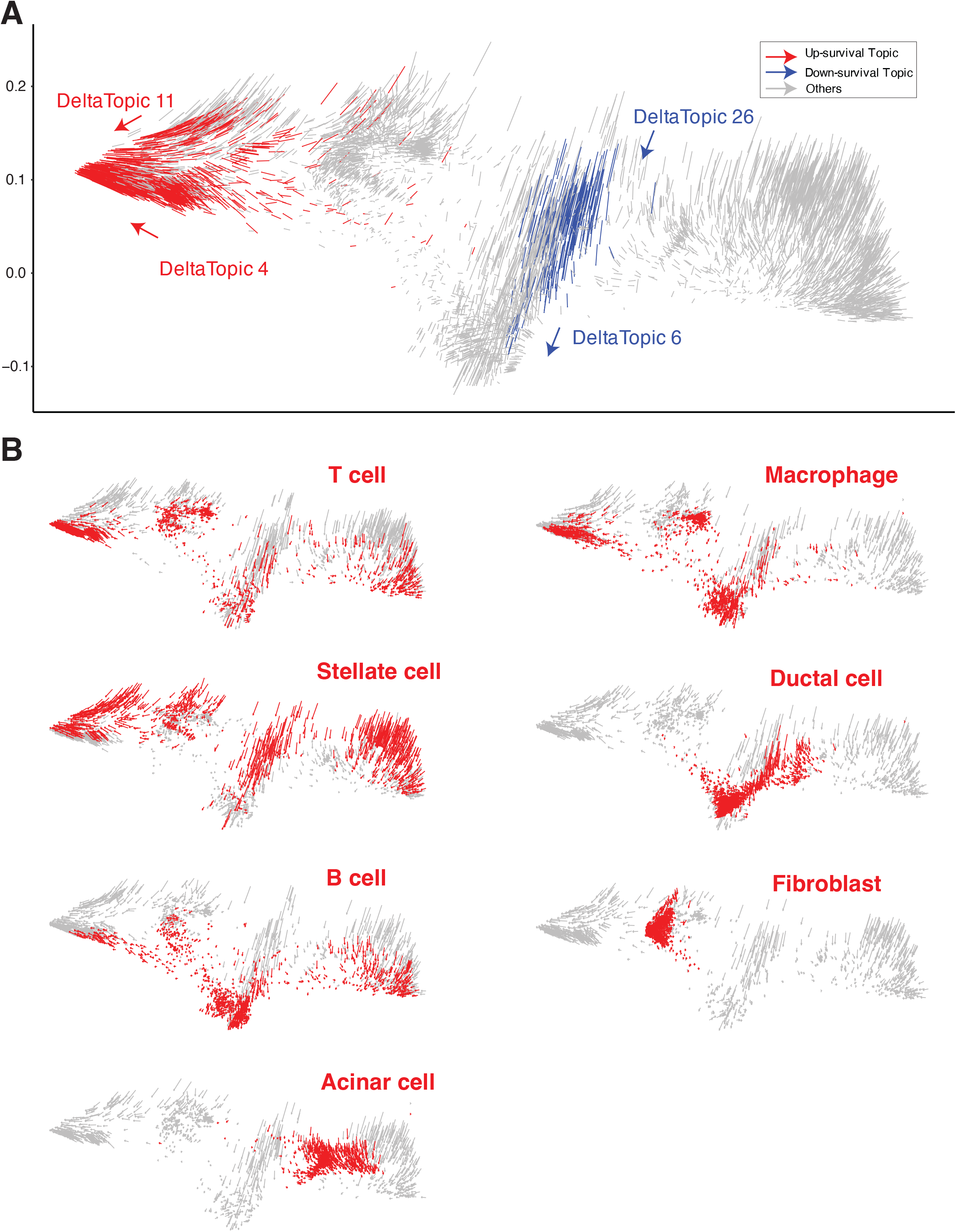
Velocities derived from the DeltaTopic identify distinct cell trajectoryies for disease development and cell type differentiation. The estimated velocities for a sample of cells. We uniformly sample cells from the population with a sample rate of 0.5% in each subfigure. **(A)** Cells in disease-relevant topics are depicted in red and blue for up and down-regulation in survival time. The up-survival and down-survival cell topics form two distinct velocity streamlines, pointing toward two directions. **(B)** Velocities are highlighted in red for seven cell types.

Disease topics generally constitute two distinct flow patterns, consistent with our previous survival analysis results. Cell topics 4 and 11, which play the up-regulation role in PDAC survival, converge to the left of the figure. On the other hand, the two down-regulating cell topics, 6 and 26, mostly flow in a downward direction (Fig. 4A). Three immune cell types (T-cell, B-cell, and macrophage) align well with this disease-specific direction. However, Acinar cells and fibroblasts are relatively stagnant, not showing any salient flows in our visualization.

## Discussion

In this study, we propose a novel Bayesian topic model built on a deep-learning-based framework, firstly incorporating well-established sparsity prior distribution to model parameters (BALSAM) and secondly incorporating short-term dynamics implicated by the difference between the spliced and unspliced reads in single-cell RNA-seq data (DeltaTopic). Considering technical limitations posed by the limited range of transcriptomic profiling, we believe that our Bayesian approach can provide a practical and statistical approach to estimating transcriptomic dynamics without requiring unattainable steady-state observations. In our case studies, we have demonstrated that DeltaTopic models built on two types of data modalities (the spliced and unspliced counts) achieved better generalization performance in terms of hold-out data Reconstruction performance. In a broader sense, our approach can be considered a special case of a mixture of generalized linear models (GLM),^30^ which augments GLMs into traditional mixture components to express conditional probabilities between two different data types.

By incorporating epigenomic and proteomic measurements, our approach can be straightforwardly extended to capture full information flows in regulatory genomics problems. Perhaps, a harder challenge still lies in linking regulatory elements to target genes and selecting isoforms to target proteins and protein complexes. Nonetheless, we argue that employing multiple layers of generalized linear models to link data modalities can provide a principled way to address more complex multiomics data integration problems. As long as multiple stages of first-order approximation can constitute full-phase diagrams of a system of differential equations, we expect such piecewise, layer-by-layer approaches will be used effectively in future research.

Interestingly, in our pancreatic cancer analysis, looking at short-term dynamics improved our understanding of disease progression. Our DeltaTopic model can pick up cancer-progression-specific latent factors, and the significance of our findings was validated in larger cohorts. Since we know relevant cell types where these disease-specific factors are selectively activated/repressed, we expect further *in vitro* or *ex vivo* validation experiments will further elucidate novel aspects of disease mechanisms. Although many existing latent variable models focus on clustering and subtype identification, our model especially highlighted short-term dynamics (merely directional information) can provide an important clue in translation studies. Tracking full trajectories of disease progression may be intractable in current technology, but predicting immediate next steps at each cell type/state level appears possible. Conversely, dynamics analysis can complement a current way of investigating cellular heterogeneity, suggesting that there are other amiss axes in disease mechanisms besides cell type composition changes.

As suggested by the previous Embedded topic modelling approach,^31^ prior knowledge can play an important role in dealing with stochastic, noisy data sets. In our case, eventually, we will need to understand transcription factors that drive topic-specific dynamics and disease mechanisms. Again, incorporating histone and DNA accessibility data will greatly benefit multimodal single-cell analysis.

## Acknowledgments

This work was supported by the BC Cancer Foundation and NSERC Discovery Grant (YPP). We are deeply grateful for support from Adrian Wan and the BC Cancer IT team.

## Author contributions

Conceptualization, YPP; Methodology, YZ, MSK, and YPP; Investigation, MSK and YZ; Writing – Original Draft, YZ and YPP; Writing – Review & Editing, YPP; Funding Acquisition, YPP; Resources, YPP; Supervision, YPP

## Figures

**Supplementary Figure 1.**

**(A)** Circular chord diagram for the matching between delta topics and Balsam topics. An edge is drawn between two topics if they are assigned to the same cell based on the loadings. Topics with less than 2% of the total population cells are removed from the chord diagram.

## Methods

### Single-cell RNA-seq data preparation

We obtained the original FASTQ files for pancreatic ductal adenocarcinoma (PDAC) from the public repository (https://ngdc.cncb.ac.cn/gsa/browse/CRA001160) provided by two PDAC studies.^6,7^ The spliced and unspliced count matrices were quantified by a scalable approach, namely `kb-python` (https://www.kallistobus.tools/), which coordinates the inputs and outputs of Kallisto^8^ and UMI-BUS^9^ tools:

~~~
$ kb count -i index.idx -g t2g.txt -x 10xv2 -o ${output} \
    -c1 spliced_t2c.txt -c2 unspliced_t2c.txt \
    --workflow lamanno --filter bustools \
    ${fastq1} ${fastq2}
~~~

### Notations

We measured *G* genes on a total of *N* cells. We denote the spliced one by *X_ng_* for a gene *g* in a cell *n*, capturing only the reads mapped on exonic regions, gene data used in most single-cell expression analyses. We use 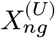 to denote an unspliced expression activity for a gene *g* and a cell *n*, concerning reads mapped on, or involving intronic regions. For brevity, we will use a row vector x_*n*_ = (*X*_*n*1_,…,*X_nG_*) for expression data (spliced count) on each cell *n*. Similarly, we will use 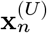 to denote an “unspliced” expression count, 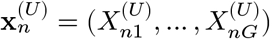.

Following the mapping protocol proposed by La Manno and coworkers,^1^ we can separately quantify 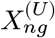, the count of unspliced transcripts of a gene *g* in a cell *n*, and 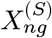, the count of the spliced of the gene *g* in the same cell *n*.

### Model descriptions

#### Review of topic modelling

Embedded Topic Model^12^ built on a Variational Auto-Encoder (VAE) framework^32^ provides a scalable approach for discovering latent topics from a large corpus of documents. We applied a similar topic model approach to single-cell data, treating each cell as a document, 20k genes as a total set of vocabularies, and short reads mapped on the genes as words. We model gene expression counts generated by independent multinomial probabilities (a bag of words assumption) since multinomial likelihood better preserves scale-invariant properties across different batches than other deep learning methods based on Poisson, Negative Binomial, and Gaussian distributions. Letting *X_ng_* be a gene expression count of a gene *g* in a cell *n*, the data likelihood of a total expression count matrix can be defined by the corresponding multinomial probabilities *r_ng_*:

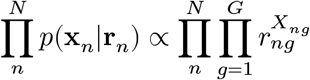

where we have ∑*_g_ r_ng_* = 1 for all *n* ∈ [*N*] and *r_ng_* > 0 for all cells *n* and genes *g* ∈ [*G*].

Dieng and coworkers proposed a powerful and scalable way to estimate these multinomial probabilities as a linear combination of topic-specific vocabulary (gene) matrices, *γ_tg_*, weighted by topic portions in a document, 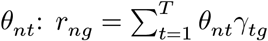.

For simplicity, the latent topic proportion vectors are assumed to follow Logistic Normal distribution *a priori*, which can enable reparameterized variational inference by taking stochastic gradient steps. We generate *θ_n_* in two steps: 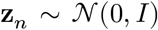 and *θ_nt_* = exp(*z_nt_*)/∑_*t*′_ exp(*z_nt′_*). Since we restrict both *θ* and *γ* in a *T*- and *G*-dimensional simplex, namely, ∑_*t*_ *θ_nt_* = 1 and ∑_*g*_ *γ_tg_*, we can confirm that the resulting **r**_*n*_ vectors also result in valid probabilities across vocabulary (genes) within cells, i.e., 0 ≤ ∑_*g*_ *r_ng_* ≤ 1.

#### A Bayesian extension for topic modelling in single-cell RNA-seq analysis

We extended the embedded topic model (ETM) in two ways: (1) We introduced a Bayesian hierarchical prior on the model parameters, **r**_*n*_ ~ Dir(*λ_n_*) while formulating the Dirichlet parameters as a generalized linear model,

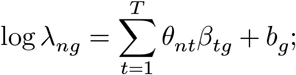

(2) We sought to improve the interpretability of the model parameters by introducing Bayesian sparsity on the linear models, *β_tg_*, assuming a majority of the *β_tg_* values are statistically zero with some prior probability *π*,

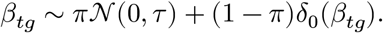

The other effects not captured by spike-and-slab *β* parameters are simply represented by a gene-specific bias parameter *b_g_* invariantly present across all the topics. We did not enforce any specific prior distribution on the bias parameters.

Exploiting the conjugate relationship between the multinomial and Dirichlet distributions, we can analytically integrate out the composite variable *r* and derive the following data likelihood:

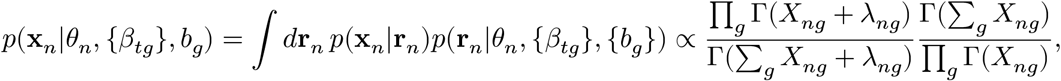

where 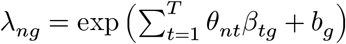, and Γ(·) is the Euler’s gamma function.

#### Amortized variational inference

As the dimensionality increases, the exact inference of posterior probability of the latent cell-specific topics p(*θ_n_*|x_*n*_) quickly turns into a computationally-intractable inference problem. Stochastic variational inference confers a scalable approach to finding approximating distributions, which often leads to surprisingly accurate posterior inference results.^33^ A reparametrization technique popularized by the VAE framework^32^ cast an intractable inference problem of a latent variable model into an optimization problem in a deep belief network model, which can be solved by taking back-propagation steps with respect to the model parameters.^34^ Here, we use two types of variational distributions, namely one for the local, latent variables, *q*(*θ_n_*) and the other for global topic-specific gene activity parameters *q*(*β_tg_*), and minimized the Kullbeck-Leibler (KL) divergence between these approximates and the actual data likelihood probability models.

Equivalently we can maximize the following evidence lower bound (ELBO) for total data likelihood (denoted by 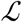):

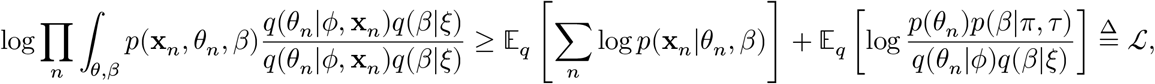

where we used *ϕ* and *ξ* to denote all the parameters of the latent state/parameter distributions. Amortized variational inference algorithm finds approximate posterior distributions by optimizing the ELBO objective, taking stochastic gradient steps with respect the variational parameters, namely *ϕ* and *xi*. We adaptively scheduled the learning rates and step sizes by Adam optimizer.^35^ For gradient calculation, we used `PyTorch` library.

##### Sampling from the latent variable model *q*(*θ|ϕ*)

Since the exact evaluation the first expectation term is generally intractable, we approximate it by summing over the data log-likelihood using the sampled instances of *θ*^(*s*)^ ~ *q*(*θ*|*ϕ*, x_*n*_) and *β*^(*s*)^ for each minibatch sample *s* ∈ [*S*]:

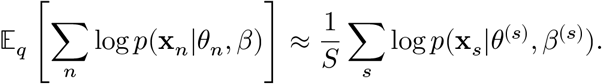

We parameterized the mean *μ* and variance *σ* functions for the latent variable inference in a deep encoder model taking inputs of the original high-dimensional data **x**. Using the reparamterized trick of Logistic Normal distribution, we sample the posterior sample of *θ*^(*s*)^ as follows:

- Sample 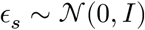
- Reparamterize *z_s_* ← *μ*(x_*s*_) + *σ*(x_*s*_) ∘ *ϵ_s_*
- Transform 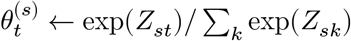.

Using the corresponding variational parameters *ϕ* ≡ (*μ, σ*), assuming 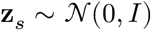 *a priori* for all *s*, we can derive KL divergence between the prior and variational distributions of latent states:

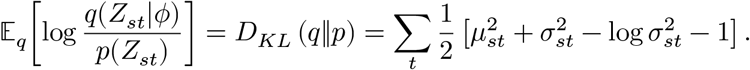

##### Global spike-and-slab parameters *q*(*β*)

We analytically derived the second term (the negative KL loss) involving the global parameters *β* by using fully-factored spike-and-slab distributions^13^ as variational distributions for *β_tg_* parameters. When *β_tg_* is active/on, more precisely, a latent indicator variable *h_tg_* = 1 with probability *α_tg_*, we parameterize it by a Gaussian distribution:

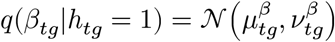

with probability 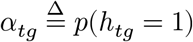; otherwise, we simply set to zero:

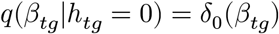

with probability 1 – *α_tg_*.

Given the variational parameters, *ξ* ≡ (*α, μ, ν*), we can characterize the mean, 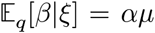 and variance, 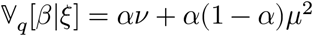.

Letting *h_tg_* = 1 with probability *π* and 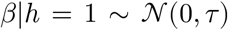 *a priori*, we get the KL loss for the global parameters:

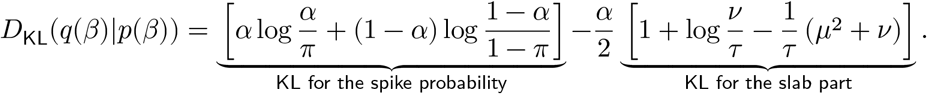

#### Dynamically-Encoded Latent Transcriptomic Analysis by Topic modelling (deltaTopic)

DeltaTopic is a hierarchical Bayesian model designed to capture transcription dynamics in topic space manifested in spliced and unspliced single-cell count matrices. Built on the Bayesian extension previously discussed, the goal of DeltaTopic is to characterize topic-specific relationships between spliced (*S*) and unspliced (*U*) gene expressions as a generalized linear model, delineating the static/shared and dynamic/directional topic-specific gene components.

We used the same Multinomial-Dirichlet hierarchical model that the unspliced and spliced vectors are parameterized by the rates of the unspliced and the unspliced, respectively:

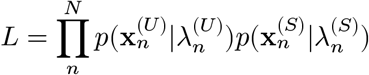

where

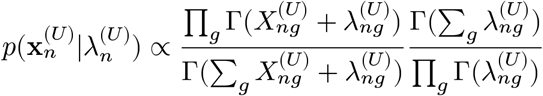

and

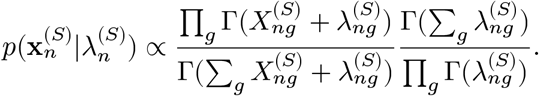

We can reasonably assume that each latent cell topic proportion vector *θ_n_* is the property of a cell *n*. In the shared topic space, the unspliced and spliced counts are differently expressed by different topic-specific rate parameters. We represent shared transcription rates (before splicing) by *ρ_tg_* of a gene *g* for a topic *t*; we explicitly capture splicing rates by *δ_tg_* of a gene *g* for a topic *t* along with a gene-specific baseline activity *b_g_*. Putting them altogether, we have the log-rates for the unspliced:

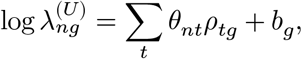

and the log-rates for the spliced:

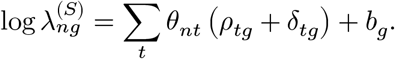

We pose spike-and-slab priors on the *ρ* and *δ* parameters:^13^

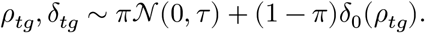

The latent vector *θ_n_* is sampled from Logistic-Normal distribution. We used two independent encoder networks and combined stochastic latent vectors as a mixture of experts, equally weighting.^14,15^ Each encoder network generates the mean *μ* and standard deviation *σ*.

- Sample 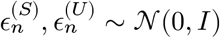
- Reparameterize for the unspliced, 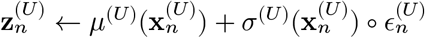
- Reparameterize for the spliced, 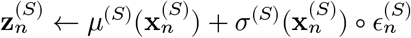
- Combine and transform: 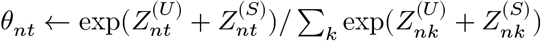.

With the above sampling scheme, we optimized the following ELBO and estimated posterior distributions of the latent states and model parameters:

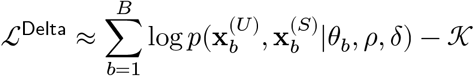

where *b* denotes an index for a mini batch sample with the batch size *B* and 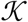 the KL loss.

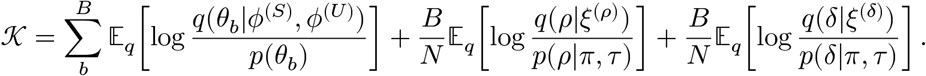

#### Kaplan-Meier survival analysis

We obtained topic-specific gene loading scores, such as *δ, rho*, and *β* parameters, after fully training topic models. We then estimated individual-level scores by multiplying them to individual-level gene expression profiles available in larger ICGC cohorts. Followed by standardization of these individual-level scores, we can stratify these individuals into the positive (Δ*_it_* > 0) and negative activity (Δ*_it_* < 0) sets for each topic t. Using the same procedure, we can also partition individuals into positively and negatively correlated groups based on the other two types of scores (*P_it_* and *B_it_*). We estimated Kapler-Miere(KM) survival curve for each topic and tested the two-group difference in survival probabilities by log-rank test.

